# Network Identification Methods

**DOI:** 10.1101/071217

**Authors:** Young Hwan Chang, Tomlin J. Clair, ICBP Member

## Abstract

Recently, network inference algorithms have grown tremendously in the field of systems biology because network identification is essential for understanding relationships between regulation mechanisms for genes, elucidating functional mechanisms underlying cellular processes, as well as identifying molecular targets for discoveries in medicines. This article provides a brief overview of different approaches used to identify biological networks and reviews recent advances in network identification.

## I. INTRODUCTION

A network is a set of nodes and a set of directed or undirected edges between the nodes. In biological systems, there exist many types of networks, including: transcriptional, signaling and metabolic networks, and an edge in the network corresponding to a biochemical interaction can be validated experimentally. However, since the size of the search space increases exponentially with the number of nodes in the network, still less is known about the structure of such networks.

Many computational methods [1–14] used to infer gene regulatory networks (GRNs) or biochemical interactions provide a prediction of the ‘wiring diagram’ of the network. Roughly, these methods identify the network structure from gene expression profile data by searching for patterns of correlation or conditional probabilities that indicate causal influence, or by finding best parameters in the mathematical model that fit the data. These approaches fall generally into the following categories: 1) Statistical models and 2) Mechanistic network models.

In general, the network identification remains a difficult problem; statistical dependencies are affected by both direct and indirect path of nodal interactions; nonlinearities in the system dynamics and measurement noise make this problem even more challenging. Also, in order to continue to have an impact in systems biology, identification of the graph topology from data should be able to reveal deficiencies in the model and suggest new experimental directions.

## II. STATISTICAL APPROACH

Statistical approaches use the so-called ‘influence’ network model, which generally reflects global properties of a system's behavior, and thus true molecular interactions are described rather implicitly [15]. Hence, these models can be difficult to interpret and also difficult to integrate further information.

### A. Correlation-Based Method

Predictions of physical and functional links between cellular components are often based on correlations between experimental measurements, such as gene expression [1]. The correlation coefficient *ρ*_*ij*_ (between gene *i* and *j*) determines the connection between gene *i* and *j*; Adjacency matrix (*A*) can be defined as follows:

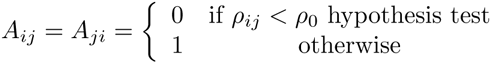

This gives an undirected, unweighted network. However, many methods relying on a variety of pairwise gene expression correlation measures are subject to exceedingly high false-positive rates (direct vs. indirect influence, i.e., via one or more intermediaries).

### B. Mutual Information (MI)

Mutual information is a measure of the mutual dependence between two variables which can be expressed by the joint distribution of two random variable X and Y relative to the joint distribution of X and Y under the assumption of independence. The mutual information can be defined as *I(X;Y)=H(X)-H(X|Y)* where *H(X)* is the marginal entropy and *H(X|Y)* is the conditional entropy. By applying the data processing inequality [16], indirect interactions can be eliminated since statistical dependencies might be of an indirect nature. For example, if *X*➔*Y*➔*Z*, then *I(X;Y) ≥ I(X;Z)*, with equality if and only if *I(X;Y|Z) =0*.

Many approaches apply some additional filtering and post-processing procedures and the final result is an adjacency matrix from which we can infer interactions.

### C. Bayesian Network

Graphical model is a term that refers to the separation of a joint probability distribution into conditional probabilities. It is commonly used in Bayesian networks, which have several attractive properties for the inference of signaling pathways from biological data sets; Bayesian networks can represent stochastic nonlinear relationships and describe direct molecular interactions as well as indirect influences that proceed through additional unobserved components. Thus, very complex relationships in signaling pathways can be discovered [17–19].

In the formulation of Bayesian networks, the structure of a genetic regulatory network is modeled by a directed acyclic graph *G* = *(V, E)* where vertices (*V*) represent genes or other elements and edges (*E*) represent biochemical interactions in the network. Bayesian network modeling associates with each variable *X*_*i*_, a probability distribution conditioned on its parents in the graph (*Pa*_*i*_). The graph structure represents the dependency assumptions that each variable is independent of its non-descendant; thus the joint distribution can be decomposed into the following form:

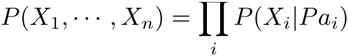

The goal of Bayesian network inference is to search among possible graphs and select the best graph which describes the dependency relationships observed in the experimental data. One can take a score-based approach and given a scoring function and a set of data, network inference amounts to finding the structure that maximizes the score. Main challenges include: exponential complexity in the local network connectivity necessitating heuristic search procedures, reliance on unrealistic network models and the need to discretize expression data [15].

## III. MECHANISTIC MODELING APPROACH

Mechanistic network models identify the interactions based on a prior knowledge being used as biologically motivated constraints, i.e., reducing search space. Thus, such reverse engineering approaches reveal the best interaction maps that fit the data to prior models [20,21]. Many methods consider a dynamical system that depends on a reaction graph, summarizing all biochemical reactions and associated parameters. These methods assume that neither the graph nor the parameters are known. Inference regarding the graph structure is carried out by integrating experimental data with dynamic models and then reformulating parameter estimation problems. In this way, one can take account of model complexity as well as the fit-to-data.

### A. Boolean Networks

The state of a gene can be described by a Boolean variable, i.e., a gene is considered to be either ‘on’ or ‘off’, and intermediate expression levels are neglected and hence its products are present or absent. Using Boolean variables, interactions between states can be represented by a Boolean functions, which define the status of state of a gene from the activation of other genes. Also, modeling regulatory networks by means of Boolean networks allows large regulatory networks to be analyzed in an efficient way, by making strong assumptions on the structure and simple dynamics of GRNs.

### B. Ordinary Differential Equations

Ordinary differential equations (ODEs), which model the dynamics of biological systems, have been widely used to analyze GRNs. The ODE formulation models the concentrations of RNAs, proteins, and other molecules by time-dependent variables with values contained in the set of nonnegative real numbers. Regulatory interactions take the form of functional and differential relations between the concentration variables. Specifically, gene regulation is modeled by rate equations expressing the rate of production of a component of the system as a function of the concentrations of other components as follows
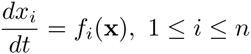

where **x** is the vector of concentrations of proteins, mRNAs, or small molecules and *f*_*i*_ represents a nonlinear function. With dynamic models, regression techniques fit the data to *a priori* model and we can infer interaction maps from the biochemical reactions and associated parameters [22].

## IV. RECENT TRENDS IN NETWORK IDENTIFICATION

Although high-throughput measurement techniques have grown tremendously, still data insufficiency strongly impedes identification of GRNs. Hence, in order to obtain reliable inference results, it is important to incorporate biologically motivated constraints (i.e., sparsity). Also, many researchers propose new methods which combine diverse types of data together (e.g., multidimensional -omic data, ChIP-on-chip data, protein-protein interaction data, sequence information), or integrate a number of independent experimental clues from literature or biological databases.

In general, regression techniques fit the data to prior models and such methods are limited to relatively simple models, i.e., usually based on simple, often linear, approximations to underlying dynamics. This is due to the fact that as the network complexity increases, the number of parameters becomes much larger than the number of experimental constraints. Thus, incorporating biologically motivated constraints is very useful to reduce the search space. Since biological regulatory networks are known to be sparse, meaning that most genes interact with only a small number of genes compared with the total number in the network, many methods take advantage of the sparsity [22]. These methods typically use *l*_*1*_-norm optimization, which leads to a sparse representation of the network and improves the ability to find the actual network structure. Moreover, these methods can be extended to combine *a priori* information on the network structure (i.e., known promotion and inhibition relations can be coded in with constraints).

Various information from scientific literature and biological database can be used in combination with experimental data. Recently, many researchers proposed promising methods that integrate such diverse types of data in GRN identification [23]. Thus, facing limited amounts of experimental data, the integration of prior biological knowledge and multiple sources of heterogeneous data will be one of the important focuses in future GRN identification research.

## Notes

This research was supported by the NIH NCI under the ICBP and PS-OC programs (5U54CA112970-08)

## REFERENCES

[1] Ideker T, Ozier O, Schwikowski B, Siegel AF: Discovering regulatory and signaling circuits in molecular interaction networks. Bioinfomatics 2002, 18:233–240.

[2] Ma L, Iglesiass PA: Quantifying robustness of biochemical network models. BMC Bioinformatics 2002, 3(38):1–13.

[3] Sontag ED: Network reconstruction based on steady-state data. Essays Biochem 2008, 45:161–176.

[4] Zechnera C, Ruessa J, Krenn P, Pelet S, Peter M, Lygeros J, Koeppl H: Moment-based inference predicts bimodality in transient gene expression. Proc Natl Acad Sci U S A 2012, 109(21):8340–8345.

[5] Zavlanos MM, Julius AA, Boyd SP, Pappas GJ: Identification of stable genetic networks using convex programming. In Proceedings of the American Control Conference (ACC). Seattle, WA: IEEE; 2008:2755–2760.

[6] Cooper NG, Belta CA, Julius AA: Genetic regulatory network identification using multivariate monotone functions. In Proceedings of the IEEE conference on Decision and Control and European Control Conference (CDC-ECC). Orlando, FL: IEEE; 2011:2208–2213.

[7] Porreca R, Drulhe S, de Jong H, Ferrari-Trecate G: Structural identification of piecewise-linear models of genetic regulatory networks. J Comput Biol 2008, 15(10):1365–1380.

[8] Bernardo DD, Gardner TS, Collins JJ: Robust identification of large genetic networks. Pac Symp Biocomput 2004, 9:486–497.

[9] Richard G, Julius A. A, Belta C: Optimizing regulation functions in gene network identification. In IEEE Conference on Decision and Control (CDC). Firenze, Italy: IEEE; 2013: 745–750.

[10] Gonçalves JM, Warnick SC: Necessary and sufficient conditions for dynamical structure reconstruction of lti networks. IEEE Trans Automatic Control 2008, 53(7):1670–1674.

[11] Sontag E, Kiyatkin A, Kholodenko BN: Inferring dynamics architecture of cellular network using time series of gene expression, protein and metabolite data. Bioinformatics 2004, 20(12):1877–1886.

[12] Han S, Yoon Y, Cho K-H: Inferring biomolecular interaction networks based on convex optimization. Comput Biol Chem 2007, 31(5-6):347–354. 14. Napoletani D, Sauer TD: Reconstructing the topology of sparsely connected dynamical networks. Phys Rev E 2008, 77(2):026103.

[13] Napoletani D, Sauer T, Struppa DC, Petricoin E, Liotta L: Augmented sparse reconstruction of protein signaling networks. J Theor Biol 2008, 255(1):40–52.

[14] Yuan Y, Stan G-B, Warnick S, Gonçalves J: Robust dynamical network structure reconstruction. Automatica 2011, 47: 1230–1235.

[15] Michael Hecker, Sandro Lambeck, Susanne Toepfer, Eugene van Someren, Reinhard Guthke, Gene regulatory network inference: Data integration in dynamic models—A review, Biosystems, Volume 96, Issue 1, April 2009, Pages 86–103

[16] Adam A Margolin, Kai Wang, Wei Keat Lim, Manjunath Kustagi, Ilya Nemenman, Andrea Califano, Reverse engineering cellular networks, Nature Protocols 1, 662–671 (2006)

[17] Sachs, K., O. Perez, D. Pe’er, D.A. Lauffenburger, and G.P. Nolan, “Causal Protein Signaling Networks Derived From Multiparameter Single-Cell Data”, Science 308: 523–529 (2005).

[18] Friedman N, Linial M, Nachman I, Pe’er D (2000) Using Bayesian networks to analyze expression data. J Comput Biol 7:601–620

[19] Hill, S.M., Lu, Y., Molina, J., Heiser, L.M., Spellman, P.T., Speed, T.P., Gray, J.W., Mills, G.B., Mukherjee, S. (2012) Bayesian Inference of Signaling Network Topology in a Cancer Cell Line. Bioinformatics, 28 (21), pp. 2804–2810.

[20] S. Han, Y. Yoon, and K.-H. Cho, Inferring biomolecular interaction networks based on convex optimization, Computational Biology and Chemistry, vol. 31, no. 5–6, pp. 347–354, Oct. 2007.

[21] Chang YH, Gray J, Tomlin C. Optimization-based inference for temporally evolving networks with applications in biology, J Comput Biol. 2012 Dec;19(12):1307–23. doi: 10.1089/cmb.2012.0190.

[22] Chang YH, Gray JW, Tomlin CJ. BMC Bioinformatics. Exact reconstruction of gene regulatory networks using compressive sensing, 2014 Dec 14;15(1):400.

[23] Young Hwan Chang, Roel Dobbe, Palak Bhushan, Joe W. Gray, Claire Tomlin, “Reconstruction of gene regulatory networks based on repairing sparse low-rank matrices”, the Seventh Annual RECOMB/ISCB RSG Conference, Nov 9–14, San Diego 2014

